# Longitudinal single cell fate of hematopoiesis *in vivo* using cellular barcoding and DiSNE movie visualization

**DOI:** 10.1101/279406

**Authors:** Jerry Gao, Dawn S. Lin, Edmund Crampin, Shalin H. Naik

**Affiliations:** Molecular Medicine, The Walter & Eliza Hall Institute of Medical Research, Parkville, Victoria 3052, Australia; Immunology, The Walter & Eliza Hall Institute of Medical Research, Parkville, Victoria 3052, Australia; Faculty of Medicine, Dentistry & Health Sciences, University of Melbourne, Parkville,Victoria 3010, Australia; Systems Biology Laboratory, University of Melbourne, Parkville, Victoria 3010, Australia; Centre for Systems Genomics, University of Melbourne, Parkville, Victoria 3010, Australia; ARC Centre of Excellence in Convergent Bio-Nano Science and Technology, Melbourne School of Engineering, University of Melbourne, Parkville, Victoria 3010, Australia

## Abstract

Identifying the progeny of many single progenitor cells simultaneously can be achieved by tagging progenitors with unique heritable DNA barcodes, and allows inferences of lineage relationships, including longitudinally. While this approach has shed new light on single cell fate heterogeneity, data interpretation remains a major challenge. In this study, we applied our developmental interpolated t-Distributed Stochastic Neighbor Embedding (DiSNE) movie approach to visualize the clonal dynamics of hematopoietic reconstitution in primates and identify novel developmental patterns, namely a potential cluster of hematopoietic progenitors with early T cell and later granulocyte production.

**Key points:**

- Complex single cell haematopoietic fate heterogeneity can be visualized and assessed with tSNE pie maps
- DiSNE movie visualization of *in vivo* haematopoiesis allows “play back” of the waves of haematopoiesis
- Identification of novel hematopoietic progenitors with early T cell and later granulocyte production

## Introduction

Hematopoiesis is a dynamic process where a given stem or progenitor cell may differ in its quantitative contribution to different lineages, and this may vary over time. Consensus models of hematopoiesis are still in flux, and contributed by population and single cell approaches through RNA-sequencing and fate analysis (Naik et al., 2013b; Paul et al., 2015; Rodriguez-Fraticelli et al., 2018; Sanjuan-Pla et al., 2013; Velten et al., 2017). One consistent feature, however, is that, stem and progenitor fractions give rise to lineages at different times(Forsberg et al., 2006). In these ‘waves’, Mk/E, myeloid and DC lineages peak early (*i.e.*, 1-2 weeks), whereas lymphoid development, which requires DNA recombination for receptor rearrangement for T cells and B cells, peaks much later (*i.e.*, 4-8 weeks). These data represent population-level snapshots but do not account for how individual clones stack to generate the ensemble average.

One strategy to unravel this complexity *in vivo* involves lineage tracing of a single cell per mouse, and has been utilised in several landmark studies (Dykstra et al.,2007; Osawa et al., 1996; Yamamoto et al., 2013). These represent informative but logistically challenging experiments, generally limited to dozens or hundreds of clones in total, and where single cells are not in competition. Another strategy to study clonal heterogeneity is the use of cellular barcoding, which allows the *in vivo* fate of individual cells within a population to be assessed in a competitive setting within one animal (Lu et al., 2011; Naik et al., 2013b; 2014). Building on the seminal studies using retroviral integration site analysis(Lemischka et al., 1986), this technology relies on differential tagging of progenitor cells with unique heritable DNA sequence tags (barcodes) so that subsequent quantification and barcode comparison between progeny cell types allows inference of lineage relationships. At its simplest level, barcodes shared between cell types infers derivation from common ancestors, whereas differing barcodes infers separate ancestors. This approach has revolutionized the assessment of cell fate heterogeneity by offering a fine-grained analysis of the quantity, quality, location and kinetics of cell fate. Several studies utilizing this technology have offered novel insights into fate and clone size heterogeneity in hematopoiesis (Gerrits et al., 2010; Lu et al., 2011; Naik et al., 2013a), breast tissue and cancer development (Eirew et al., 2015; Nguyen et al., 2014a; 2014b) and T cell immunity (Gerlach et al., 2013; Schepers et al., 2008; van Heijst et al., 2009). Limited studies have further dissected fate heterogeneity through time in transplantation settings, either in mice (Verovskaya et al., 2013) or primates (Kim et al., 2014; Wu et al., 2014). Add to this the emergence of *in situ* barcode labeling techniques that read out native haematopoiesis (Höfer et al., 2016; Pei et al., 2017; Rodriguez-Fraticelli et al., 2018; Sun et al., 2014), and one can appreciate that single cell approaches are only beginning to reveal the true complexity of haematopoiesis and other developmental systems.

The data derived from cellular barcoding experiments can be complex, and interpretation remains a major challenge. Inspired by recent analyses of cytometry data using t-SNE(Amir et al., 2013; Becher et al., 2014; Van der Maaten and Hinton, 2008), we customized this algorithm for dynamic visualization and extrapolation of fate biases over time (Lin et al., 2018). Here, using this approach, we re-classify CD34^+^ hematopoietic progenitors in primates into groups of lineage-restricted progenitors that contribute in different waves in addition to stable multi-lineage contributing clones. The analysis identified a group of progenitors displaying unexpected early T cell followed by later granulocyte production.

## Materials and Methods

### Data sets and processing

Data from the Naik *et al.* (Naik et al., 2013a) and Wu *et al.* (Wu et al., 2014) studies were first bioinformatically processed and filtered as described originally(Naik et al., 2013a). Each sample was then normalized to 100,000 read counts and transformed using hyperbolic arcsin. Barcodes with no counts for all samples were removed, and this data was then used to generate t-SNE maps.

### t-SNE

MATLAB source code for t-SNE was obtained from http://lvdmaaten.github.io/tsne/ and was run using parameters as described in Amir *et al.* (Amir et al., 2013). PCA was not used as a preprocessing step prior to performing t-SNE. When investigating cell type-specific reconstitution kinetics, barcodes with no counts in the cell type of interest at any time point were removed, even if counts were present in other cell types at some time point. Also, a small, random perturbation between 0 and 0.001 was added to the barcode read counts as too many barcodes had exactly the same read count profile, which affects the clustering ability of t-SNE. When generating the dynamic t-SNE pie map movies, all progenitors were assumed to have no output to any cell type at the initial frame (time = 0 months). At intermediate frames between time points where experimental data was gathered, the clone size and relative proportions of output to the progeny cell types for each progenitor was approximated using linear interpolation.

### Manual classification on t-SNE

Manual classification of t-SNE maps was conducted such that all identified classes at least have a common dominant feature (*e.g.*, all have B cell progeny), if not the exact same qualitative output. Small classes with fewer than 10 progenitors were also avoided. Automated methods for classifying data such as *k*-means or DensVM (Becher et al., 2014) are not suitable for cellular barcoding data as they establish a centroid for each class and group data points within some degree of variation from that centroid. Other methods such as DBSCAN can be used, but were not utilized here.

### JS divergence

For computing the JS divergence, a bin size of 5 in each t-SNE dimension was used to discretize the t-SNE map. JS divergence was calculated using the *PieMaker* software package (Lin et al., 2018) (https://data.mendeley.com/datasets/9mkz5n9jtf/1).

### DiSNE movie visualization and JS divergence

DiSNE movies were generated using the *PieMaker* software package following the instruction in the user manual(Lin et al., 2018) (https://data.mendeley.com/datasets/9mkz5n9jtf/1).

### Additional algorithms

All hierarchical clustering performed in this study used Euclidian distance and complete linkage. PCA, Isomap and LLE were run using the MATLAB Toolbox for Dimensionality Reduction (van der Maaten et al., 2009) downloaded from http://lvdmaaten.github.io/drtoolbox/. For Isomap and LLE, the smallest connectivity parameter that provided a mapping for all high-dimensional data points was used.

## Results

### SNE is an effective tool for visualizing cell fate heterogeneity in mouse LMPPs

In order to first test t-SNE based visualization of multi-lineage data derived *in vivo* (Lin et al., 2018), we first evaluated heterogeneity in cell output of LMPPs in mice. We pooled previously generated cellular barcoding data(Naik et al., 2013a) from six mice across three experiments and, in order to directly compare between experiments, the six cell types that were assessable for all mice were included: from the bone marrow (BM) B cells, neutrophils and monocytes; and from the spleen cDC1, pDC and B cells. Application of t-SNE to this data created a two-dimensional map on which each point represented a barcoded LMPP, and where LMPPs with a similar output of progeny lay close together (Figure 1a). Points on the t-SNE map were then manually classified such that the qualitative (cell types produced) output of progenitors in each class was approximately the same, with at least the same dominant feature (see Supplemental Experimental Procedures for manual versus automated classification). We identified eleven classes of LMPPs using this approach (Figure 1a) and generated heat maps to visualize the clonal qualitative and quantitative (abundance of cells) output from each class (Figure 1b). The t-SNE algorithm allowed effective identification of seven mono-outcome classes - one for each cell type as well as an additional splenic cDC1-only low-output class. The remaining multi-outcome classes consisted of 1) dominant BM B cell production plus splenic B cells or cDC1s and pDCs; 2) splenic B cells and cDC1s and pDCs; 3) splenic cDC1s and pDCs; and 4) BM monocytes and neutrophils and splenic cDC1s and pDCs, with a small subpopulation of these progenitors contributing greatly to splenic B cells as well. These results showed that the t-SNE algorithm is effective in positioning the LMPPs based on their qualitative and quantitative outputs.

**Figure 1.**
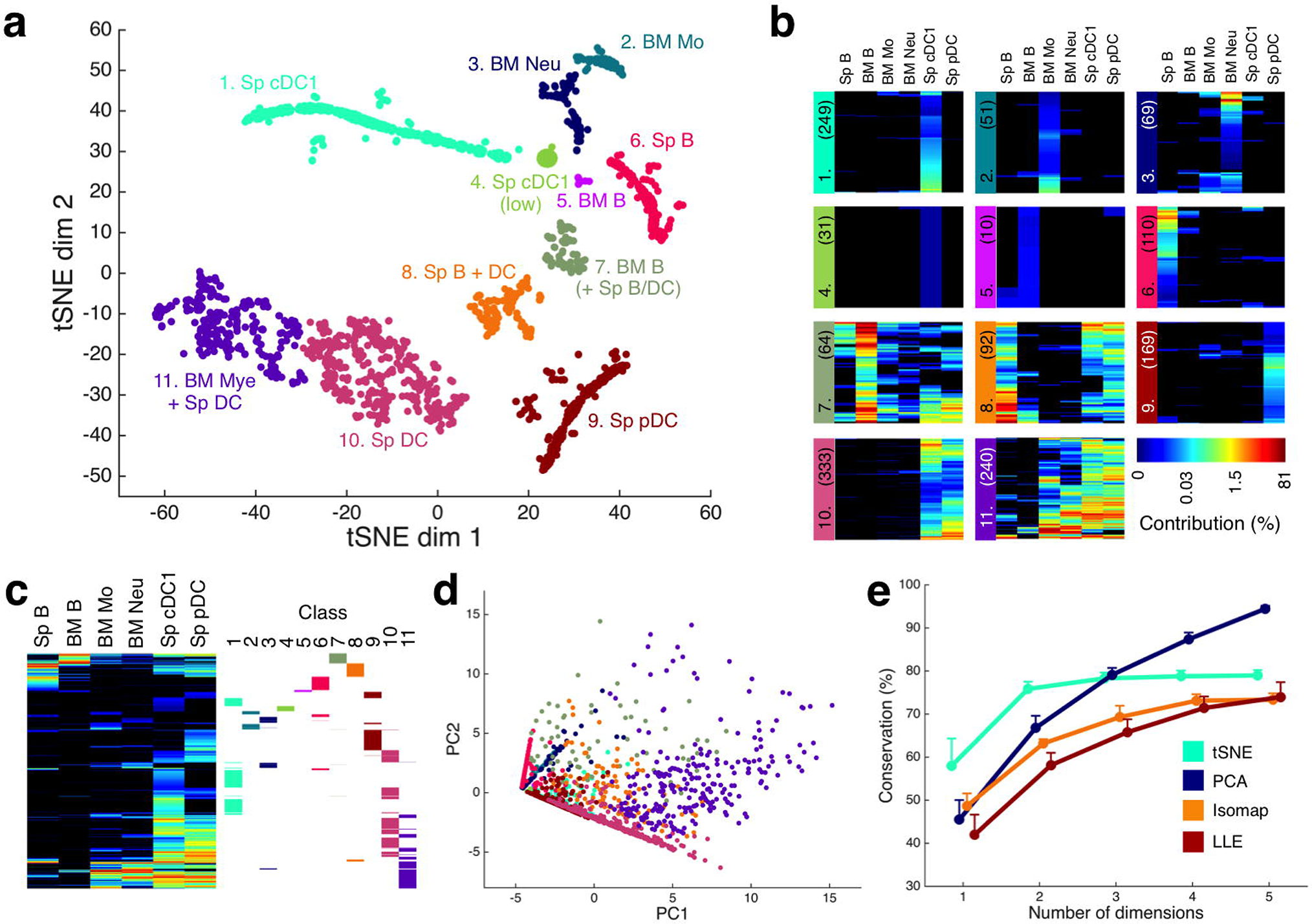
t-SNE for visualization and classification of cellular barcoding data. **a)** t-SNE map of the output of 1,418 LMPPs from six mice to six different cell types. Progenitors were manually classified and named based on their patterns of output in b. **b)** Heat maps of each class where each row represents a barcode and each column the cell type for which the barcode contributes, and color is indicative of the relative contribution of each LMPP to each cell type. Number of barcodes in each class is shown in brackets. **c)** Hierarchical clustering of the same data, with the progenitors in each t-SNE class “backgated” and highlighted in columns to the right. Note that t-SNE classes are largely dispersed down the clustered heat map, and not grouped, indicating that hierarchical clustering does not capture the same classes as t-SNE. **d)** t-SNE classes backgated onto a two-dimensional PCA of the same data. Note how the t-SNE classes are not easily separable. **e)** Comparison of different dimensionality reduction techniques for their conservation of the 20 nearest neighbors. Error bars are standard errors of the mean across the six mice. t-SNE outperforms PCA, Isomap and LLE for conservation of nearest neighbors at lower numbers of dimensions (*i.e.*, groups of progenitors with similar fate patterns are best segregated with t-SNE when plotted in two dimensions).

We then compared where the progenitors from each t-SNE class were positioned relative to the ordering determined by hierarchical clustering as shown in Figure 1c. Importantly, the classes did not always segregate, indicating that patterns observed from t-SNE analysis may not be identifiable from hierarchical clustering. To examine any advantage of t-SNE compared to classic dimensionality reduction techniques, we also “backgated” t-SNE classes onto the first two dimensions using Principal Component Analysis (PCA). t-SNE was superior to PCA in distributing LMPPs in two dimensions by visual assessment (Figure 1d). We then performed a quantitative assessment of four different dimensionality reduction techniques including t-SNE, PCA, Isomap and locally linear embedding (LLE). We identified the 20 nearest neighbors based on Euclidian distance for each LMPP and computed the proportion of nearest neighbors conserved after various degrees of dimensionality reduction (Figure 1e). t-SNE consistently outperformed Isomap and LLE, and also had a higher conservation than PCA when reducing to three dimensions or less. Considering the difficulty in visualizing information beyond two to three dimensions, t-SNE better captured groups of LMPPs with a similar fate than did any of the classical techniques, demonstrating its utility as a discovery tool for pattern identification.

Although t-SNE generated a low-dimensional representation of the data to enable intuitive exploration, further inspection was necessary to infer meaning. To facilitate this, we applied a recently developed visualization tool called the PieMaker {Lin:2018gk} and generated “t-SNE pie map” for individuadl clones. We scaled point size according to the clone size of that progenitor (*i.e.*, the summed output of each LMPP across all progeny cell types), and each point was also replaced by a pie chart portraying the relative proportions of output to the progeny cell types (Figure 2a). From this visual representation, we noticed that almost all LMPPs from the mono-outcome classes only contributed a small amount to their respective cell type, and conversely, within the multi-outcome classes existed individual LMPPs that had major contribution to one or more cell types (Figure 2b). The “t-SNE pie map” thereby facilitated visualization of both cell fate and clone size heterogeneity, and revealed potential positive correlation between the two clonal properties.

**Figure 2.**
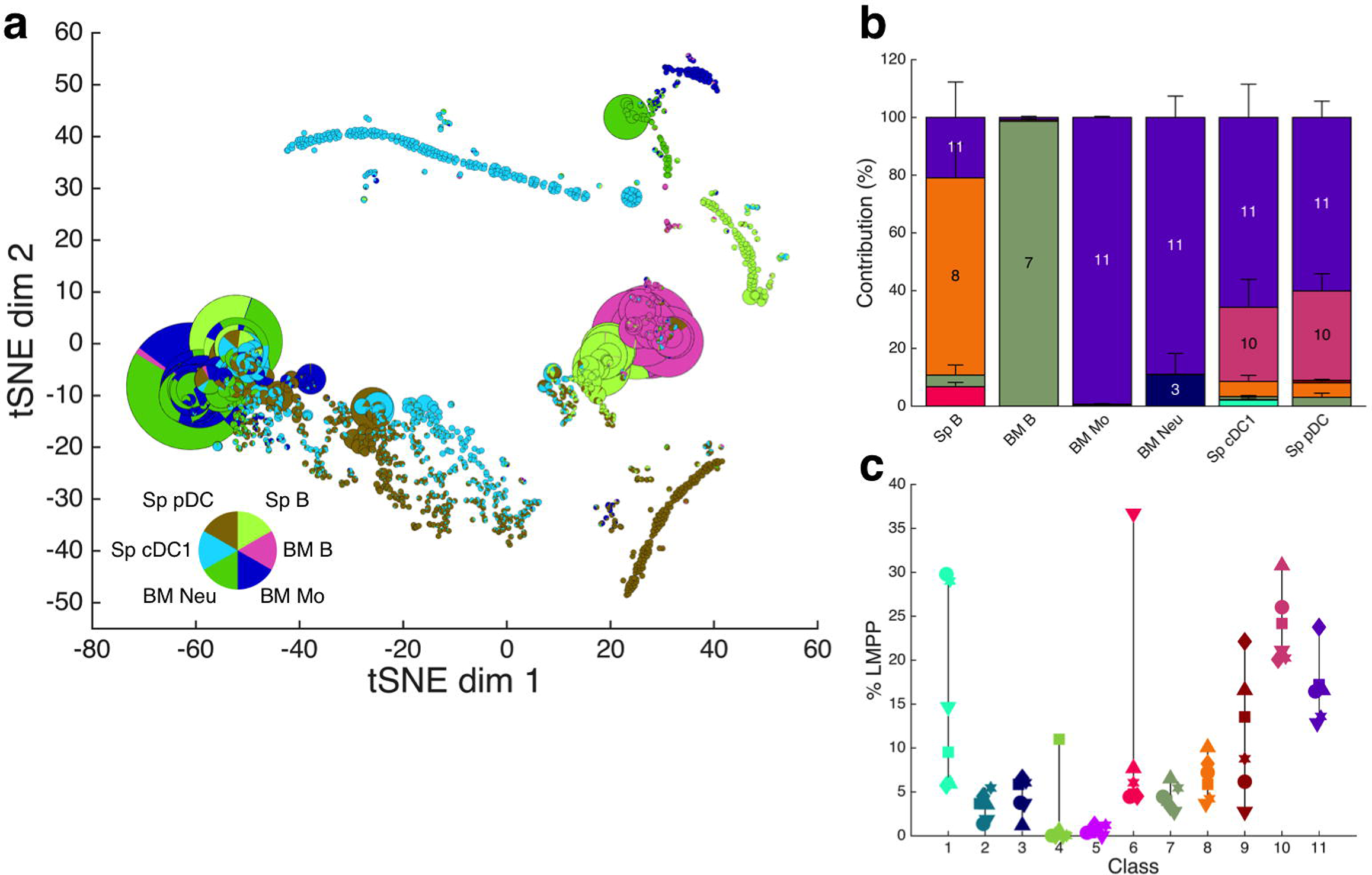
Behavior and reproducibility of LMPP classes identified using t-SNE. **a)** t-SNE pie map of the same data as in Figure 1a, with point size scaled according to the clonal output of each progenitor, and converted to a pie chart to visualize the relative contribution of that clone to the different cell types. **b)** Contribution of t-SNE classes to each cell type, with classes contributing to 10% or more to a cell type labeled. Error bars are standard errors of the mean across the six mice, and class numbering and color is consistent with Figure 1. The dominantly contributing classes are all multi-outcome. **c)** The proportion of LMPPs from each of six mice present in each class. Lines span range of LMPP proportions for each class, and each symbol represents an individual mouse. Most classes have representative LMPPs from each mouse.

A major challenge in analyzing single cell tracking data is to assess biological reproducibility of any identified patterns of cell fate, and applying t-SNE to the pooled data of six mice allowed such assessment. By overlaying mouse identity onto all LMPPs, we observed that most classes contained representatives from each mouse (Figure S1a). We also quantified this as a proportion of LMPPs per mouse in each class (Figure 2c) and found that most classes showed similar representation levels across mice, indicating reproducibility of those classes. In some classes, such as class four (low output cDC1-only progenitors) and six (splenic B cell-only progenitors), there was high representation in a single mouse. Whether this reflected a *bona fide* class of LMPPs that was not reproducibly detected due to suboptimal sampling, a difference in the environment of that mouse, or instead a spurious event, could be determined by examining more mice or a greater number of progenitors per mouse. Therefore, as described previously{Lin:2018gk}, t-SNE not only allows identification of major patterns of cell fate, but also assessment of biological reproducibility.

The Jenson-Shannon (JS) divergence can be applied to t-SNE maps as a holistic and quantitative ‘similarity’ metric (Amir et al., 2013). The minimum divergence of 0 indicates exactly the same distribution of progenitors on the t-SNE map, whereas the maximum divergence of 1 indicates no overlap. To assess animal variability, we first calculated the JS divergence for each pair of mice and, following hierarchical clustering, found that mice from the same experiment were similar with JS divergence <0.5 (Figure S1b). In fact, mice one to four from experiments one and two consistently had a pairwise divergence <0.5, but mice from experiment three (especially mouse five) showed greater differences from the other mice. We also computed JS divergences for each mouse against the other mice pooled (*e.g.*, mouse one against mice two to six pooled), and found that the divergences were substantially lowered. This indicated that a sampling effect may have contributed to the large fluctuations in divergences between pairs of mice, and increasing the number of progenitors from each mouse could decrease this variability between animals. This strategy of applying t-SNE on pooled data followed by computing the JS divergence facilitates a quantitative comparison between samples, and could be used to compare the effect of different conditions to stem and progenitor fate (*e.g.*, host animals in the steady-state versus inflammation).

### Hematopoietic progenitors from primates consist of lineage-restricted sequential contributors and multi-lineage stable clones

To visualize longitudinal single cell tracking data, we customized a recently developed framework, which included our developmental-interpolated t-SNE (DiSNE) movie and spindle plot visualization of clusters (Lin et al., 2018). We obtained data a recent study (Wu et al., 2014), where hematopoietic stem and progenitor cells from three rhesus macaques were barcode-labeled and their output to peripheral blood T cells, B cells, natural killer (NK) cells, monocytes and granulocytes were tracked over time. Cellular output was assessed at 1, 2, 3, 4.5, 6.5 and 9.5 months, with the final time differing for each animal: 6.5, 9.5 and 4.5 months for rhesus macaques ZH17, ZH33 and ZG66 respectively.

We first inspected the degree of variation in progenitors between the different rhesus macaques by generating a single t-SNE map for all progenitors that contributed at any time point from the three animals (Figure S2a). Points on the t-SNE map again represented single progenitors but which now clustered according not only to a similar pattern of output, but also dependent on their kinetics of contribution. Note that in pooling data across the three rhesus macaques, the last time point incorporated was 4.5 months, the time up to which data was available for all animals. Surprisingly, although progenitors from ZH33 and ZG66 fell in similar regions of the t-SNE map (JS divergence: 0.38), the majority of the ZH17 progenitors lay outside these regions (JS divergence: 0.88 and 0.90 compared with ZH33 and ZG66 respectively, and 0.88 compared with ZH33 and ZG66 pooled), indicating very different reconstitution kinetics in ZH17 compared to the other two animals. We therefore proceeded with analyzing individual animals without pooling data.

To generate DiSNE movies, we performed t-SNE on data from each rhesus macaque separately, incorporating all time points available for that animal, and the resulting t-SNE maps were converted into pie maps each focusing on a single time point (Figure 3 and S2b-c). Although the entire data set across all time points was embedded within each individual pie, each of these only displayed the behavior of progenitors at its respective time point, without capturing the dynamic component. We then linked these series of static t-SNE pie maps between time points to generate DiSNE movies for dynamic visualization of the reconstitution process encompassing the complexities of qualitative, quantitative, and temporal (order and timing of contribution) characteristics of individual clones (Movies S1-3).

**Figure 3.**
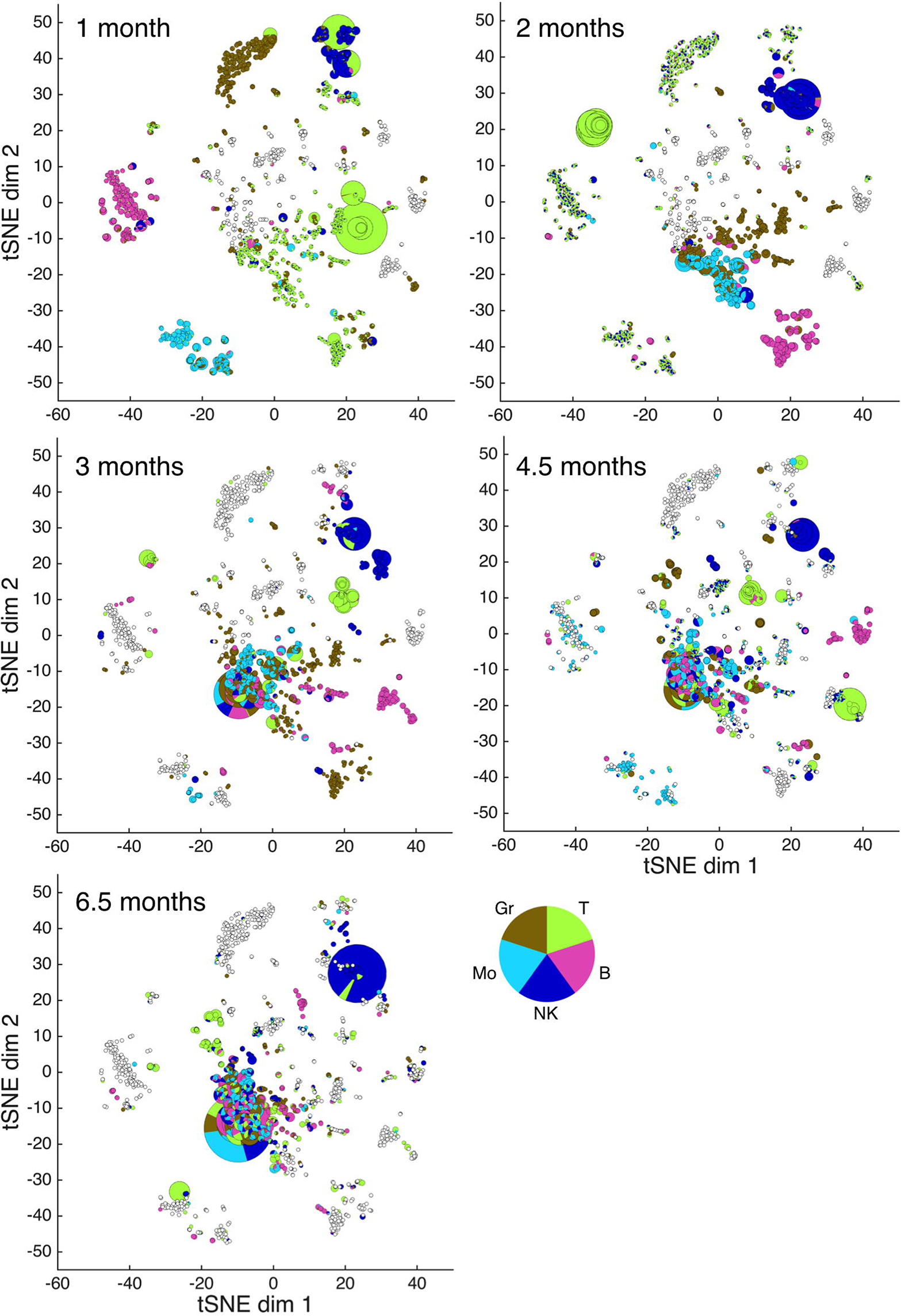
Static visualization of reconstitution kinetics in ZH17. t-SNE pie maps of the output from 1,354 hematopoietic stem and progenitor cells from ZH17 to five different cell types sampled at five time points. The underlying t-SNE map (where each progenitor is located in the reduced t-SNE dimensions) is constructed from the entire data set (*i.e.*, all cell types at all time points), but the point size and pie chart proportions for each panel are set according to the respective specific time point. Empty (white) points indicate no contribution at that time point. See Movie S1 for dynamic visualization. Lineage-restricted sequential contributors are primarily located along the periphery of the map and are the main contributors at earlier time points, and later on their activity is replaced by the multi-lineage stable clones positioned in the centre of the map. Note the persistence of the NK-specific progenitors in the top right corner of the map through all time points sampled.

When comparing the DiSNE movies and static t-SNE pie maps between individual animals, we noticed distinct patterns in ZH17 (Movie S1 & Figure 3) compared to ZH33 (Movie S2 & Figure S2b) and ZG66 (Movie S3 & Figure S2c). In ZH17, we observed very pronounced waves of transient contribution from progenitors with progeny limited to a single dominant cell type (Figure 3). This lineage-restricted sequential contribution was particularly evident in the lymphoid lineages, with transient T and B cell clones dominating production for up to 4.5 months. Although myeloid-restricted sequential contribution was also observed at 1 month, myeloid production by 2 months was stably associated with multi-lineage clones. On the other hand, lineage-restricted sequential contribution was much less obvious in ZH33 and ZG66, with stable multi-lineage clones as the major contributors as early as 2 months (Figure S2b-c). Importantly, a class of stable NK cell-restricted progenitors that persisted up to the last time point was strikingly apparent in the t-SNE pie maps from all animals, supporting the main finding from the original study (Wu et al., 2014) that NK cells have a distinct lineage origin (Figure 3 and S2b-c).

To uncouple the production of each individual cell type from the derivation of others to appreciate ‘per lineage’ reconstitution kinetics, we performed t-SNE on data for each cell type separately, incorporating all time points available (Figure 4a showed t-SNE on B cell output). These t-SNE maps were then manually classified, and the contribution of each class to that cell type through time was visualized in a spindle plot (Figure 4b for B cells and S3 for all cell types). Examining individual cell types separately again demonstrated two main progenitor classes: transient sequential, versus stable contribution, with a higher representation of sequential clones in ZH17 compared to the two other animals, which were reconstituted with stable, multi-lineage clones early in time. As seen in Figure S3, there was a large variety of patterns in which clones contributed to hematopoiesis. Particularly interesting were progenitors that contributed at multiple non-consecutive time points (*e.g.*, class 15 of B cells in ZH17), indicating a termination of contribution followed by subsequent re-contribution.

**Figure 4.**
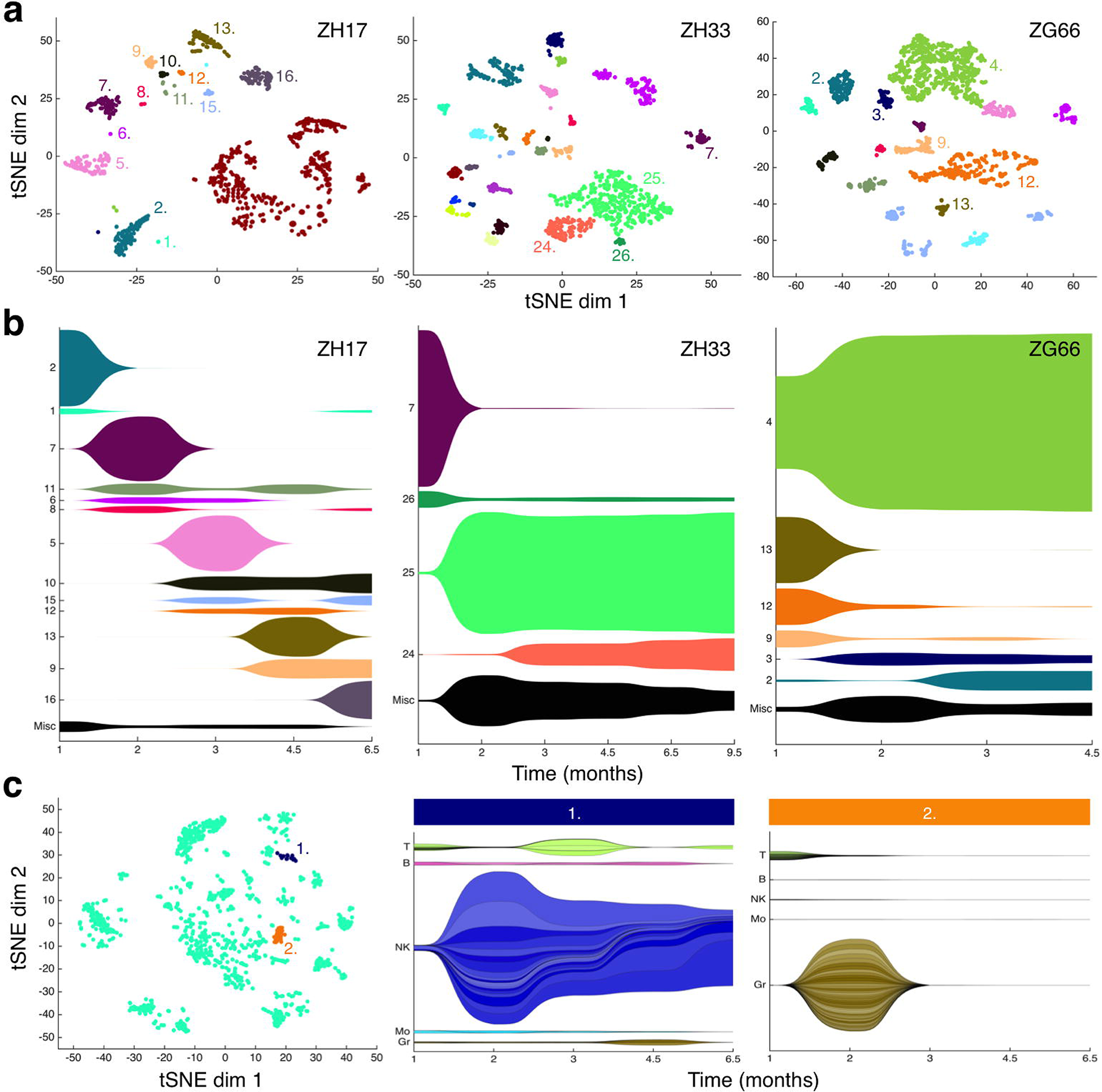
Reconstitution kinetics identified using t-SNE. **a)** t-SNE maps of the B cell output of progenitors from the three rhesus macaques across all time points sampled. Progenitors were manually classified, and those that contributed to 5% or more of all B cells at any time point are labeled. **b)** Spindle plots depicting the patterns of B cell reconstitution in the three rhesus macaques. Each spindle represents a class from a (except Misc, see below), and the width of the spindle indicates the amount of B cell contribution of that class at that time point. Classes which do not contribute to 5% or more of all B cells at any time point are not shown individually, but are instead all agglomerated and plotted as the miscellaneous (Misc) spindle at the bottom of each panel. Reconstitution kinetics in ZH17 showed very pronounced waves of transient contribution, whereas stable clones played a much greater role in ZH33 and ZG66. Note that each cell type of each animal was analyzed independently and hence class numbers across cell types and animals are also independent. **c)** Two groups of progenitors from ZH17 are highlighted on the t-SNE map considering all cell types (left panel), and the activities of progenitors from these groups are visualized using spindle plots (middle and right panels). The width of each spindle indicates the amount of contribution of the group of progenitors to the corresponding cell type at that time point, and each partition of the spindle represents an individual progenitor within the group. Partitions of the spindles are shaded randomly for ease of visualization. Group 1 primarily contributes stably to NK cells, whereas group 2 predominantly contributes transiently to granulocytes at 2 months. Note that although a few progenitors from group 1 contribute to cell types other than NK cells at various time points, all progenitors from both groups contribute to T cells at 1 month. This is especially interesting in group 2 as granulocyte production is thought to precede T cell production.

Using spindle plots, we also highlighted the activities considering all cell types of two particular groups of progenitors from ZH17 through time (Figure 4c). In one group we focused on the progenitors contributing stably to NK cells, reiterating the main finding of study (Wu et al., 2014) in an visually intuitive manner. In another group, we identified progenitors initially producing T cells followed later by granulocyte production. This ordering of events is peculiar as granulocyte production is thought to precede that of T cells, and co-production in the absence of other cell types is rare. As this cluster of progenitors was only identified in one primate, further studies should confirm or exclude the existence of progenitors with such behavior and the mechanisms responsible. Our method therefore facilitated classification, which when presented as spindle plots, intuitively captured the complexity and dynamics of hematopoietic reconstitution.

## Discussion

The analysis of high-dimensional data is a broad challenge in biology, and a major factor hampering progress is our inability to ‘see’ the data. By applying our recently developed DiSNE visualization tools we were able to identify and assess, *de novo*, patterns of reproducibility in *in vivo* models of haematopoiesis in mice and primates. In particular, we identified rare subsets with unique size, fate and temporal properties. This is exemplified by the identification of a subset of CD34^+^ hematopoietic progenitors that had an early T cell (without B cell) production wave, with a later granulocyte (without monocyte) production wave (Figure 4c, population 2). This represents a subset of progenitors that was missed in the original macaque barcoding study (Wu et al., 2014). It is novel both for the unexpected order of development (T cell development typically proceeds granulocyte production (Forsberg et al., 2006; Luc et al., 2008; Reya et al., 2001)) and that no other cell types within their respective lineages were co-produced. This is the first report of such a progenitor and, while such a pattern would need to be validated further, demonstrates the power of our computational approach for novel pattern identification.

This finding also fits with a growing body of evidence of unusual pairings of fate from stem/progenitors (Ceredig et al., 2009), including for T cell development with other myeloid cells (Bell and Bhandoola, 2008; Bhandoola et al., 2007; De Obaldia et al., 2013). However, as these studies tend to study output at the population level, and at a single time point, our findings add fine-grained temporal and clonal complexity in understanding the dynamics of hematopoiesis. This data is consistent with a ‘graded’ or ‘probabilistic’ model where commitment can occur at all stages of hematopoiesis, *i.e.*, currently defined progenitor populations contain a mixed population of committed in-transit cells, and uncommitted cells (Naik, 2008; Nimmo et al., 2015). A longer-term challenge will be to fit these data into a revised model that is more informative and accurate than current oversimplified stick-and-ball bifurcation models of hematopoiesis.

Another challenge will be identifying the causative factors of fate heterogeneity whether they are intrinsic or extrinsic. Emerging high throughput studies using single cell RNA-sequencing are revealing such regulators, however definitive linking of each gene to its bona fide fate profile is precluded due to the destructive nature of scRNA-seq. Such knowledge may have clinical benefit if, for example, one could isolate or engineer the sub-fraction of CD34^+^ cells that represent true-long term stable HSCs for transplantation, or select for transient progenitors that generate a restricted repertoire of cell types. We are currently exploring experimental means to explore and link fate heterogeneity with gene expression heterogeneity to uncover fate programs at the single cell level. The method described herein provides the framework to rapidly examine such relationships and appreciate biological processes at a systems level.

## Acknowledgments

The authors wish to thank Dr. Samir Taoudi and Prof. Doug Hilton for insightful discussions and comments on the manuscript.

This work was funded by GNT1062820 from the National Health & Medical Research Council, Australian Government.

## Authorship Contributions

JG performed the analysis, developed *PieMaker* and wrote the manuscript. DSL wrote the manuscript. EC provided critical support and guidance, and commented on the manuscript. SHN conceived the study, performed the biological interpretation and wrote the manuscript.

## References

Amir, E.-A.D., Davis, K.L., Tadmor, M.D., Simonds, E.F., Levine, J.H., Bendall, S.C., Shenfeld, D.K., Krishnaswamy, S., Nolan, G.P., Pe’er, D., 2013. viSNE enables visualization of high dimensional single-cell data and reveals phenotypic heterogeneity of leukemia. Nature Biotechnology 31, 545–552. doi:10.1038/nbt.2594

Becher, B., Schlitzer, A., Chen, J., Mair, F., Sumatoh, H.R., Teng, K.W.W., Low, D., Ruedl, C., Riccardi-Castagnoli, P., Poidinger, M., Greter, M., Ginhoux, F., Newell, E.W., 2014. High-dimensional analysis of the murine myeloid cell system. Nat. Immunol. 15, 1181–1189. doi:10.1038/ni.3006

Bell, J.J., Bhandoola, A., 2008. The earliest thymic progenitors for T cells possess myeloid lineage potential. Nature 452, 764–767. doi:10.1038/nature06840

Bhandoola, A., Boehmer von, H., Petrie, H.T., Zuniga-Pflucker, J.C., 2007. Commitment and Developmental Potential of Extrathymic and Intrathymic T Cell Precursors: Plenty to Choose from. Immunity 26, 678–689. doi:10.1016/j.immuni.2007.05.009

Ceredig, R., Rolink, A.G., Brown, G., 2009. Models of haematopoiesis: seeing the wood for the trees. Nature Publishing Group 9, 293–300. doi:10.1038/nri2525

De Obaldia, M.E., Bell, J.J., Bhandoola, A., 2013. Early T-cell progenitors are the major granulocyte precursors in the adult mouse thymus. Blood 121, 64–71. doi:10.1182/blood-2012-08-451773

Dykstra, B., Kent, D., Bowie, M., McCaffrey, L., Hamilton, M., Lyons, K., Lee, S.-J., Brinkman, R., Eaves, C., 2007. Long-Term Propagation of Distinct Hematopoietic Differentiation Programs In Vivo. Cell Stem Cell 1, 218–229. doi:10.1016/j.stem.2007.05.015

Eirew, P., Steif, A., Khattra, J., Ha, G., Yap, D., Farahani, H., Gelmon, K., Chia, S., Mar, C., Wan, A., Laks, E., Biele, J., Shumansky, K., Rosner, J., McPherson, A., Nielsen, C., Roth, A.J.L., Lefebvre, C., Bashashati, A., de Souza, C., Siu, C., Aniba, R., Brimhall, J., Oloumi, A., Osako, T., Bruna, A., Sandoval, J.L., Algara, T., Greenwood, W., Leung, K., Cheng, H., Xue, H., Wang, Y., Lin, D., Mungall, A.J., Moore, R., Zhao, Y., Lorette, J., Nguyen, L., Huntsman, D., Eaves, C.J., Hansen, C., Marra, M.A., Caldas, C., Shah, S.P., Aparicio, S., 2015. Dynamics of genomic clones in breast cancer patient xenografts at single-cell resolution. Nature 518, 422–426. doi:10.1038/nature13952

Forsberg, E.C., Serwold, T., Kogan, S., Weissman, I.L., Passegue, E., 2006. New evidence supporting megakaryocyte-erythrocyte potential of flk2/flt3+ multipotent hematopoietic progenitors. Cell 126, 415–426. doi:10.1016/j.cell.2006.06.037

Gerlach, C., Rohr, J.C., Perie, L., van Rooij, N., van Heijst, J.W.J., Velds, A., Urbanus, J., Naik, S.H., Jacobs, H., Beltman, J.B., de Boer, R.J., Schumacher, T.N.M., 2013. Heterogeneous Differentiation Patterns of Individual CD8+ T Cells. Science 340, 635–639. doi:10.1126/science.1235487

Gerrits, A., Dykstra, B., Kalmykowa, O.J., Klauke, K., Verovskaya, E., Broekhuis, M.J.C., de Haan, G., Bystrykh, L.V., 2010. Cellular barcoding tool for clonal analysis in the hematopoietic system. Blood 115, 2610–2618. doi:10.1182/blood-2009-06-229757

Höfer, T., Busch, K., Klapproth, K., Rodewald, H.-R., 2016. Fate Mapping and Quantitation of Hematopoiesis In Vivo. Annu. Rev. Immunol. 34, 449–478. doi:10.1146/annurev-immunol-032414-112019

Kim, S., Kim, N., Presson, A.P., Metzger, M.E., Bonifacino, A.C., Sehl, M., Chow, S.A., Crooks, G.M., Dunbar, C.E., An, D.S., Donahue, R.E., Chen, I.S.Y., 2014. Dynamics of HSPC Repopulation in Nonhuman Primates Revealed by a Decade-Long Clonal-Tracking Study. Cell Stem Cell 14, 473–485. doi:10.1016/j.stem.2013.12.012

Lemischka, I.R., Raulet, D.H., Mulligan, R.C., 1986. Developmental potential and dynamic behavior of hematopoietic stem cells. Cell 45, 917–927.

Lin, D.S., Kan, A., Gao, J., Crampin, E.J., Hodgkin, P.D., Naik, S.H., 2018. DiSNE Movie Visualization and Assessment of Clonal Kinetics Reveal Multiple Trajectories of Dendritic Cell Development. Cell Reports 22, 2557–2566. doi:10.1016/j.celrep.2018.02.046

Lu, R., Neff, N.F., Quake, S.R., Weissman, I.L., 2011. tracking single hematopoietic stem cells in vivo using high-throughput sequencing in conjunction with viral genetic barcoding. Nature Biotechnology 29, 928–933. doi:10.1038/nbt.1977

Luc, S., Buza-Vidas, N., Jacobsen, S.E.W., 2008. Delineating the cellular pathways of hematopoietic lineage commitment. Seminars in Immunology 20, 213–220. doi:10.1016/j.smim.2008.07.005

Naik, S.H., 2008. Demystifying the development of dendritic cell subtypes, a little. Immunol. Cell Biol. 86, 439–452. doi:10.1038/icb.2008.28

Naik, S.H., Perie, L., Swart, E., Gerlach, C., van Rooij, N., de Boer, R.J., Schumacher, T.N., 2013a. Diverse and heritable lineage imprinting of early haematopoietic progenitors. Nature 496, 229–232. doi:10.1038/nature12013

Naik, S.H., Perié, L., Swart, E., Gerlach, C., van Rooij, N., de Boer, R.J., Schumacher, T.N., 2013b. Diverse and heritable lineage imprinting of early haematopoietic progenitors. Nature 496, 229–232. doi:10.1038/nature12013

Naik, S.H., Schumacher, T.N., Perié, L., 2014. Cellular barcoding: A technical appraisal. Experimental Hematology 42, 598–608. doi:10.1016/j.exphem.2014.05.003

Nguyen, L.V., Cox, C.L., Eirew, P., Knapp, D.J.H.F., Pellacani, D., Kannan, N., Carles, A., Moksa, M., Balani, S., Shah, S., Hirst, M., Aparicio, S., Eaves, C.J., 2014a. DNA barcoding reveals diverse growth kinetics of human breast tumour subclones in serially passaged xenografts. Nature communications 5, 5871. doi:10.1038/ncomms6871

Nguyen, L.V., Makarem, M., Carles, A., Moksa, M., Kannan, N., Pandoh, P., Eirew, P., Osako, T., Kardel, M., Cheung, A.M., Kennedy, W., Tse, K., Zeng, T., Zhao, Y., Humphries, R.K., Aparicio, S., Eaves, C.J., Hirst, M., 2014b. Clonal analysis via barcoding reveals diverse growth and differentiation of transplanted mouse and human mammary stem cells. Cell Stem Cell 14, 253–263. doi:10.1016/j.stem.2013.12.011

Nimmo, R.A., May, G.E., Enver, T., 2015. Primed and ready: understanding lineage commitment through single cell analysis. Trends in cell biology. doi:10.1016/j.tcb.2015.04.004

Osawa, M., Hanada, K., Hamada, H., Nakauchi, H., 1996. Long-term lymphohematopoietic reconstitution by a single CD34-low/negative hematopoietic stem cell. Science 273, 242–245.

Paul, F., Arkin, Y., Giladi, A., Jaitin, D.A., Kenigsberg, E., Keren-Shaul, H., Winter, D., Lara-Astiaso, D., Gury, M., Weiner, A., David, E., Cohen, N., Lauridsen, F.K.B., Haas, S., Schlitzer, A., Mildner, A., Ginhoux, F., Jung, S., Trumpp, A., Porse, B.T., Tanay, A., Amit, I., 2015. Transcriptional Heterogeneity and Lineage Commitment in Myeloid Progenitors. Cell 163, 1663–1677. doi:10.1016/j.cell.2015.11.013

Pei, W., Feyerabend, T.B., Rössler, J., Wang, X., Postrach, D., Busch, K., Rode, I., Klapproth, K., Dietlein, N., Quedenau, C., Chen, W., Sauer, S., Wolf, S., Höfer, T., Rodewald, H.-R., 2017. Polylox barcoding reveals haematopoietic stem cell fates realized in vivo. Nature 548, 456–460. doi:10.1038/nature23653

Reya, T., Morrison, S.J., Clarke, M.F., Weissman, I.L., 2001. Stem cells, cancer, and cancer stem cells. Nature 414, 105–111. doi:10.1038/35102167

Rodriguez-Fraticelli, A.E., Wolock, S.L., Weinreb, C.S., Panero, R., Patel, S.H., Jankovic, M., Sun, J., Calogero, R.A., Klein, A.M., Camargo, F.D., 2018. Clonal analysis of lineage fate in native haematopoiesis. Nature 553, 212–216. doi:10.1038/nature25168

Sanjuan-Pla, A., Macaulay, I.C., Jensen, C.T., Woll, P.S., Luis, T.C., Mead, A., Moore, S., Carella, C., Matsuoka, S., Jones, T.B., Chowdhury, O., Stenson, L., Lutteropp, M., Green, J.C., Facchini, R., Boukarabila, H., Grover, A., Gambardella, A., Thongjuea, S., Carrelha, J., Tarrant, P., Atkinson, D., Clark, S.A., Nerlov, C., Jacobsen, S.E., 2013. Platelet-biased stem cells reside at the apex of the haematopoietic stem-cell hierarchy. Nature 502, 232–236. doi:10.1038/nature12495

Schepers, K., Swart, E., van Heijst, J.W., Gerlach, C., Castrucci, M., Sie, D., Heimerikx, M., Velds, A., Kerkhoven, R.M., Arens, R., Schumacher, T.N., 2008. Dissecting T cell lineage relationships by cellular barcoding. J. Exp. Med. 205, 2309–2318.

Sun, J., Ramos, A., Chapman, B., Johnnidis, J.B., Le, L., Ho, Y.-J., Klein, A., Hofmann, O., Camargo, F.D., 2014. Clonal dynamics of native haematopoiesis. Nature. doi:10.1038/nature13824

Van der Maaten, L., Hinton, G., 2008. Visualizing data using t-SNE. The Journal of Machine Learning Research.

van der Maaten, L.J., Postma, E.O., van den Herik, H.J., 2009. Dimensionality reduction: A comparative review. Journal of Machine Learning Research 10, 66–71.

van Heijst, J.W.J., Gerlach, C., Swart, E., Sie, D., Nunes-Alves, C., Kerkhoven, R.M., Arens, R., Correia-Neves, M., Schepers, K., Schumacher, T.N.M., 2009. Recruitment of Antigen-Specific CD8+ T Cells in Response to Infection Is Markedly Efficient. Science 325, 1265–1269. doi:10.1126/science.1175455

Velten, L., Haas, S.F., Raffel, S., Blaszkiewicz, S., Islam, S., Hennig, B.P., Hirche, C., Lutz, C., Buss, E.C., Nowak, D., Boch, T., Hofmann, W.-K., Ho, A.D., Huber, W., Trumpp, A., Essers, M.A.G., Steinmetz, L.M., 2017. Human haematopoietic stem cell lineage commitment is a continuous process. Nat Cell Biol 19, 271–281. doi:10.1038/ncb3493

Verovskaya, E., Broekhuis, M.J., Zwart, E., Ritsema, M., van Os, R., de Haan, G., Bystrykh, L.V., 2013. Heterogeneity of young and aged murine hematopoietic stem cells revealed by quantitative clonal analysis using cellular barcoding. Blood 122, 523–532. doi:10.1182/blood-2013-01-481135

Wu, C., Li, B., Lu, R., Koelle, S.J., Yang, Y., Jares, A., Krouse, A.E., Metzger, M., Liang, F., Loré, K., Wu, C.O., Donahue, R.E., Chen, I.S.Y., Weissman, I., Dunbar, C.E., 2014. Clonal tracking of rhesus macaque hematopoiesis highlights a distinct lineage origin for natural killer cells. Cell Stem Cell 14, 486–499. doi:10.1016/j.stem.2014.01.020

Yamamoto, R., Morita, Y., Ooehara, J., Hamanaka, S., Onodera, M., Rudolph, K.L., Ema, H., Nakauchi, H., 2013. Clonal analysis unveils self-renewing lineage-restricted progenitors generated directly from hematopoietic stem cells. Cell 154, 1112–1126. doi:10.1016/j.cell.2013.08.007

